# Interplay between *TERT* promoter mutations and methylation culminates in chromatin accessibility and *TERT* expression

**DOI:** 10.1101/859892

**Authors:** Catarina Salgado, Celine Roelse, Rogier Nell, Nelleke Gruis, Remco van Doorn, Pieter van der Velden

## Abstract

The *telomerase reverse transcriptase* (*TERT*) gene is responsible for telomere maintenance in germline and stem cells, and is re-expressed in 90% of human cancers. Contrary to common concepts, CpG methylation in the *TERT* promoter (*TERT*p), was correlated with *TERT* mRNA expression. Furthermore, two hotspot mutations in *TERT*p, dubbed C228T and C250T, have been revealed to assist binding of transcription factor ETS/TCF and subsequent *TERT* expression. This study aimed to elucidate the combined contribution of epigenetic (promoter methylation and higher-order chromatin structure) and genetic (promoter mutations) mechanisms in regulating *TERT* gene expression in healthy skin and in melanoma cell lines (n=61). We unexpectedly observed that the methylation of *TERT*p was as high in a subset of healthy skin cells, mainly keratinocytes, as in cutaneous melanoma cell lines. In spite of the high promoter methylation fraction in wild-type (WT) samples, *TERT* mRNA was only expressed in the melanoma cell lines with high methylation or intermediate methylation in combination with *TERT* mutations. *TERT*p methylation was positively correlated with chromatin accessibility and expression in 8 melanoma cell lines. Cooperation between epigenetic and genetic mechanisms were best observed in heterozygous mutant cell lines as chromosome accessibility preferentially concerned the mutant allele. Combined, these results suggest a complex model in which *TERT* expression requires either a widely open chromatin state throughout the promoter in *TERT*p-WT samples due to high methylation or a combination of moderate methylation fraction/chromatin accessibility in the presence of the C228T/C250T mutations.

**Author summary:** PvdV and RvD formulated research goals and aims and supervised the overall progress. Wet-lab experiments, preparation of the manuscript and statistical analysis were performed by CS and CR. CS designed the novel assays. RN was involved in the experimental setup. RvD, NG and PvdV were responsible for funding acquisition. CR, RN, NG, RvD and PvdV critically reviewed the manuscript.

## Introduction

Approximately 90% of all human cancers share a transcriptional alteration: reactivation of the telomerase reverse transcriptase (TERT) gene [1, 2]. TERT encodes the catalytic subunit of the ribonucleoprotein telomerase and is capable of extending the repetitive, non-coding DNA sequence on terminal ends of chromosomes, the telomeres. As the single-stranded 5’ ends of chromosomes are shortened with each cellular division, telomeres prevent loss of coding chromosomal DNA [3-6]. Telomerase is only transcribed in a subset of stem cells in growing or renewing tissues, but through reactivation of telomerase expression, cells can extend telomeres or prevent telomeres shrinkage. This is termed telomere maintenance, which is one of the hallmarks of cancer, and allows subsequent indefinite proliferation and immortalization [3, 6-8].

Since the *MYC* oncogene has firstly been identified to activate telomerase, a variety of epigenetic or genetic mechanisms in the gene body or *TERT* promoter (*TERT*p) have followed, such as CpG methylation, histone modifications, mutations, germline genetic variations, structural variations, DNA amplification or chromosomal rearrangements [3, 5, 7].

A widely investigated mechanism that could induce *TERT* reactivation is the presence of mutations in the gene promoter [7, 9]. Horn and Huang *et al.* identified two mutually exclusive *TERT*p point mutations that are correlated to *TERT* mRNA expression by creating binding motifs for the transcription factor E26 transformation-specific/ternary complex factor (ETS/TCF) [7, 9]. These mutations, chr5:1,295,228 C>T (−124 bp from the transcription start site) and chr5:1,295,250 C>T in hg19 (−146 bp from TSS), henceforth respectively dubbed C228T and C250T, were first identified in melanoma. Furthermore, these mutations showed high prevalence in and were correlated with poor prognosis of cutaneous melanomas [4, 5, 10-12].

An additional mechanism by which a gene can be made accessible to transcription factors, facilitating gene expression, is hypomethylation of promoter CpG islands, a hallmark of euchromatin [13, 14]. Methylation located in the gene body, however, shows a positive correlation with active gene expression [15]. In stark contrast to most genes, *TERT*p hypermethylation may also allow gene expression since transcriptional repressors rely on unmethylated promoter CpGs, such as CCCTC-binding factor (CTCF)/cohesin complex or MAZ [16-18]. As such, in combination with transcription factor binding, dissociation of the repressor may result in *TERT* expression [3, 16, 19, 20]. Castelo-Branco *et al.* proposed that methylation of a specific CpG site in *TERT*p, cg11625005 (position 1,295,737 in hg19) was associated with paediatric brain tumours progression and poor prognosis [20]. This finding was later supported by the study from Barthel *et al.*, in which the CpG methylation was found to be correlated with *TERT* expression in samples lacking somatic *TERT* alterations and to be generally absent in normal samples adjacent to tumour tissue [3].

Chromatin organisation, its plasticity and dynamics at *TERT*p region have been reported as relevant players in regulation of gene expression by influencing the binding of transcription factors [21, 22]. Cancer cells are positively selected to escape the native repressive chromatin environment in order to allow *TERT* transcription [23].

In the present study, we aim to elucidate the interaction of genetic and epigenetic mechanisms in regulation of *TERT*p. We approach this by using novel droplet digital PCR (ddPCR)-based assays [24]. Human-derived benign skin cells (keratinocytes, dermal fibroblasts, melanocytes, skin biopsy samples and naevi) and melanoma cell lines were analyzed. The *TERT*p mutational status was assessed along with the absolute presence of methylation in the *TERT*p at a CpG-specific resolution. The effect of chromatin accessibility in *TERT* expression was evaluated in a subset of cultured melanoma cell lines.

## Results

### NGS-based deep bisulfite sequencing and development of a ddPCR assay to assess *TERT*p methylation fraction

We first aimed to quantitatively measure the *TERT*p methylation at a CpG-specific resolution in primary skin samples and melanoma cell lines. DNA of 44 primary skin biopsy samples and melanoma cell lines was bisulfite-converted (BC) and analysed using NGS-based deep bisulfite sequencing to assess the methylation fraction (MF) in a region of *TERT*p encompassing 31 CpG sites. The *TERT*p MF was high in some healthy skin samples, such as normal skin (∼30%), naevi (∼30%) and cultured keratinocytes (∼50%). In the latter group, in fact, the MF was as high as in cutaneous melanoma cell lines (Fig 1).

**Fig 1.**
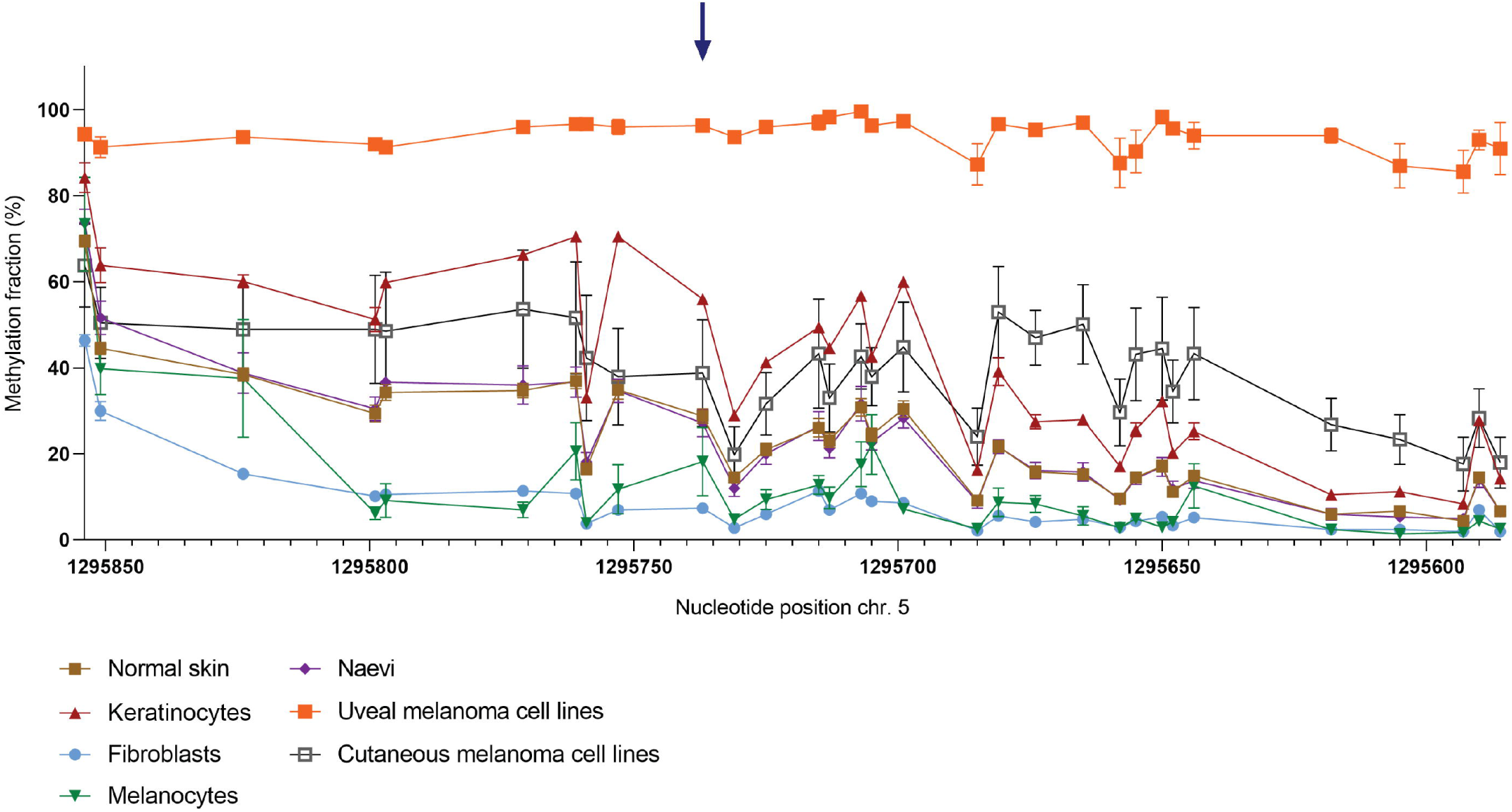
Methylation fraction (MF) of 31 CpG sites around cg11625005 in 35 primary skin samples and 9 cutaneous and uveal melanoma cell lines. DNA samples were bisulfite-converted (BC) and analysed through NGS-based deep sequencing. Connected scatter plot representing the MF per cell type group in absolute distance between measured CpG sites. Blue arrow: cg11625005 (position 1,295,737).

In order to validate the *TERT*p MF obtained through NGS in a quantitative manner, we have developed a ddPCR assay using methylation-sensitive restriction enzymes (MSREs) HgaI and AvaI, which recognise the CpG on position 1,295,737 (cg11625005) and 1,295,731 in hg19, respectively (Fig 2). Castelo-Branco *et al.* showed that methylation of the cg11625005 in *TERT*p, was associated with tumour progression and poor prognosis of childhood brain tumours [20]. Barthel *et al.* affirmed a correlation between methylation and *TERT* expression in samples lacking somatic *TERT* alterations and a lower methylation level in normal samples [3]. Indeed, in our study, the MF of fibroblasts was as low as that of the unmethylated control DNA, whereas that of the keratinocytes was higher than most of the cutaneous melanoma cell lines (Fig 2B). The MF of cg11625005 (position 1,295,737) obtained through NGS and by ddPCR were highly correlated (R^2^=0.8166, P < 0.001) (Fig 2C). The MF of 1,295,731 assessed through ddPCR even yielded a stronger correlation (R^2^=0.9580, P < 0.001) (Fig 2D).

**Fig 2.**
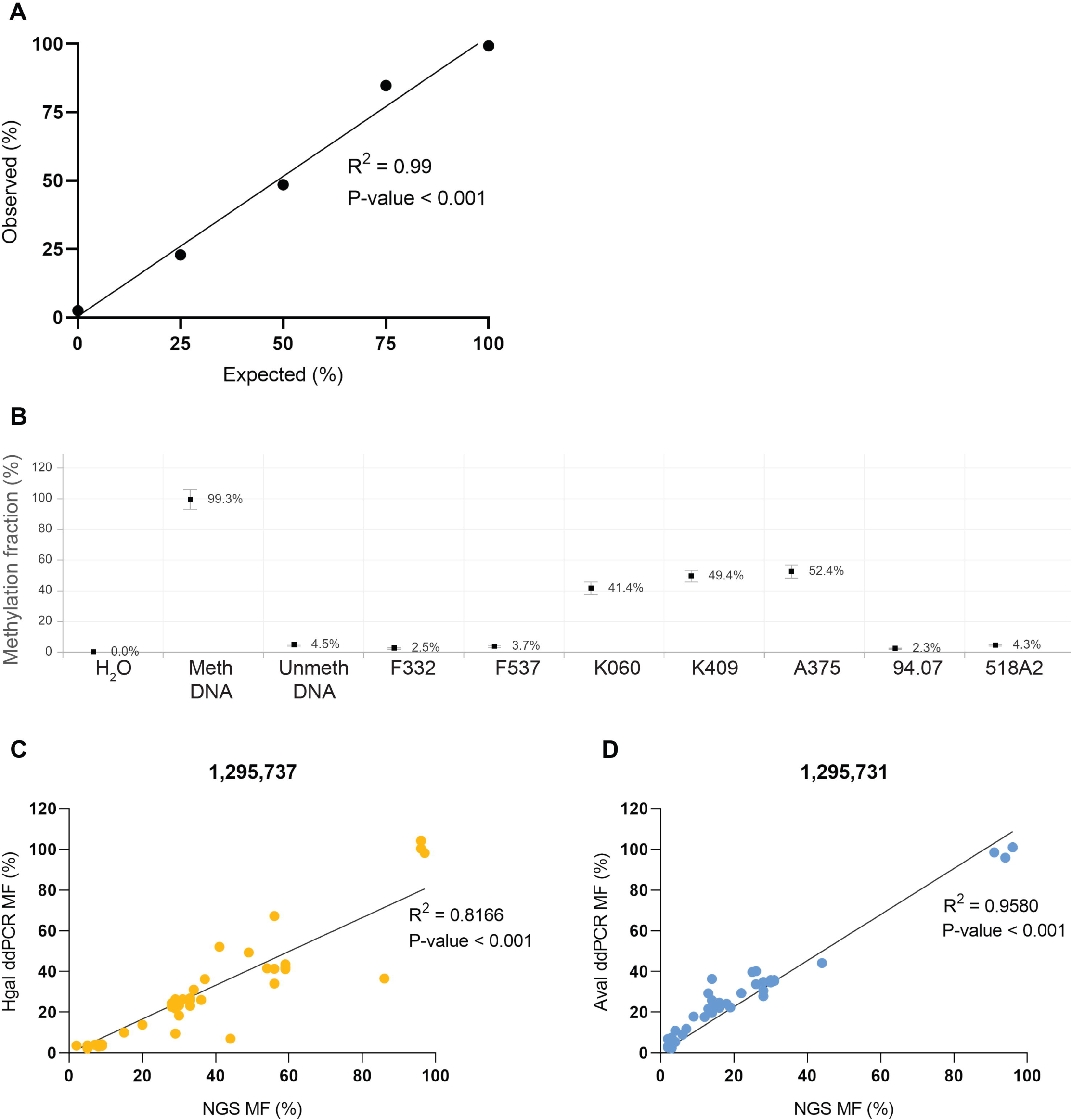
Methylation fraction (MF) analysed through ddPCR. MF was plotted through RoodCom WebAnalysis (version 1.4.2., Rogier J. Nell, Leiden). MDNA and UDNA are commercially available methylated and unmethylated DNA. **A.** Calibration curve using different expected ratios (25%, 50% and 75%) of methylated DNA and F332 to demonstrate the quantitative capacity of ddPCR. **B.** MF of cg11625005 in a subset of healthy primary skin samples – fibroblasts (F332 and F537) and keratinocytes (K060 and K409) and cutaneous melanoma cell lines (A375, 94.07 and 518A2) incubated with MSRE HgaI. **C & D**. Correlation plots between MF obtained through golden standard NGS-based deep bisulfite sequencing versus ddPCR using either the MSRE HgaI (**C.**) or AvaI (**D.**), which digest unmethylated CpG in position 1,295,737 and 1,295,731, respectively.

### Absence of correlation between methylation fraction and *TERT* expression

Cancer cells are commonly characterised by hypermethylation of promoter CpG islands resulting in repression of tumour suppressor genes. However, in *TERT*, promoter hypermethylation was found to be associated with higher expression, since CTCF repressors of *TERT* transcription do not bind methylated sequences [3, 16, 17, 19]. In our sample cohort, there was no correlation between *TERT* methylation of cg11625005 and mRNA expression (n=34, Fig 3 and an overview in Fig 7C).

**Fig 3.**
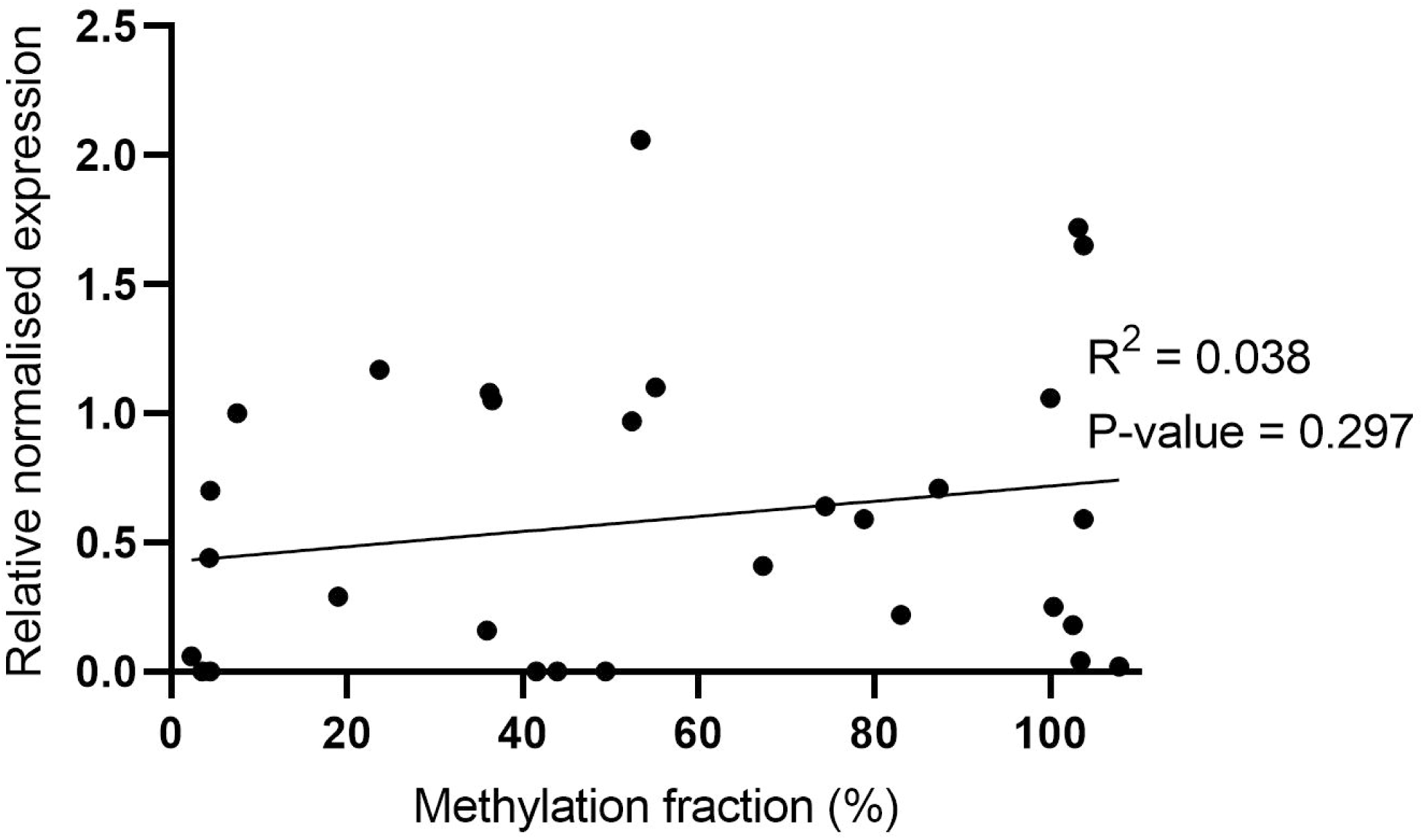
Correlation between methylation fraction (%) and *TERT* mRNA expression in total of 31 samples.

**Fig 4.**
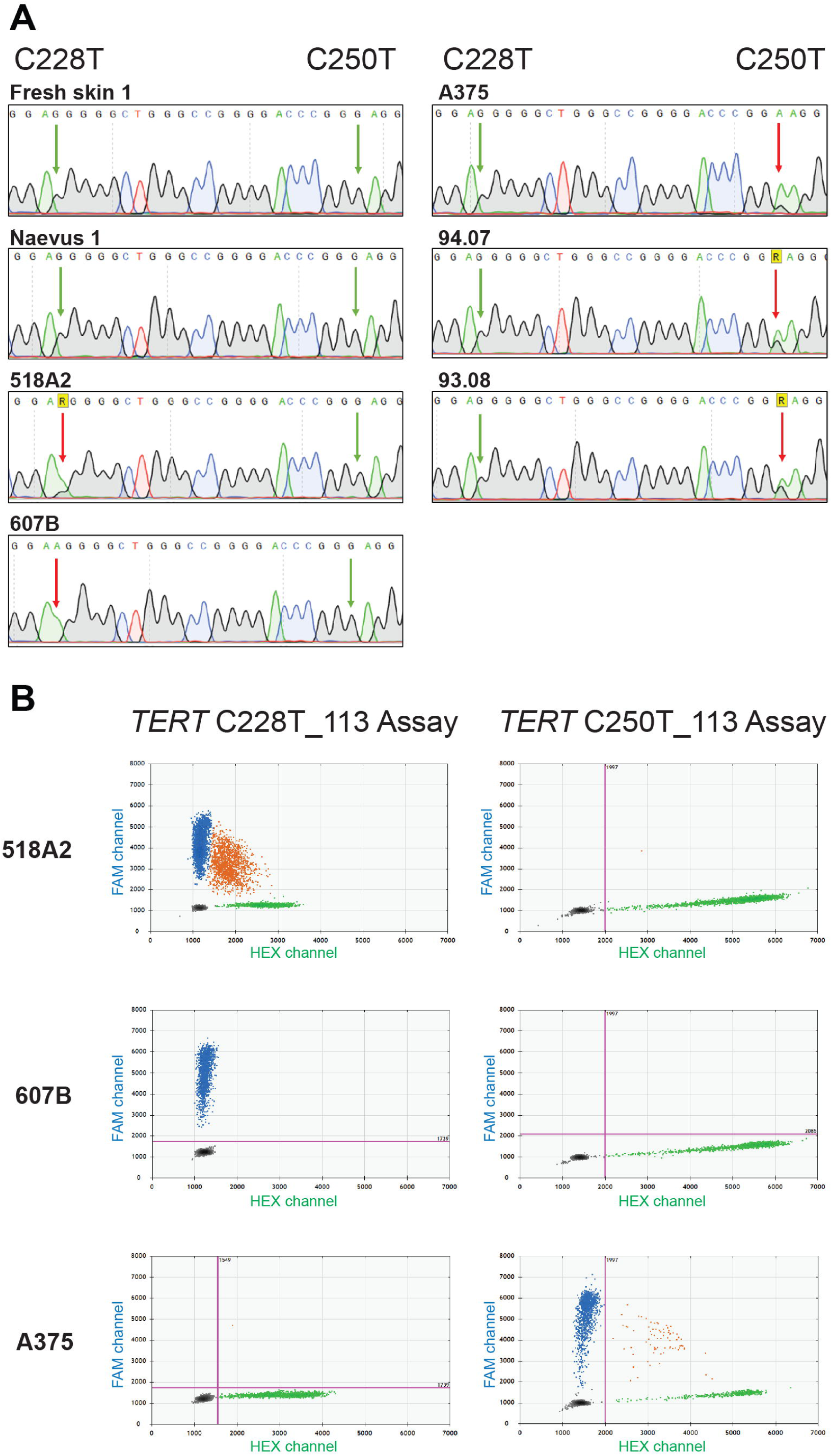
*TERT*p mutational status of primary skin samples and cutaneous melanoma cell lines. **A.** The *TERT*p region encompassing the C228T and C250T mutations was sequenced through Sanger sequencing using McEvoy’s (McEvoy et al., 2017) *TERT*p forward primer. The *TERT*p region of fresh skin 1, Naevus 1, 518A2, 607B, A375, 94.07, 93.08 is shown. The left and right arrows respectively indicate the positions 1,295,228 and 1,295,250. R: one-letter code for bases G or A; Green arrow: wild-type; red arrow: C>T mutation on the complementary strand. **B.** Evaluation of *TERT*p mutations through commercial Bio-Rad TERT assays in 518A2, 607B and A375 melanoma cell lines. 2D ddPCR plots of the results from the C228T mutation assay (left) and C250T mutation assay (right). The blue cloud represents mutant copies; the green cloud represents WT copies.

**Fig 5.**
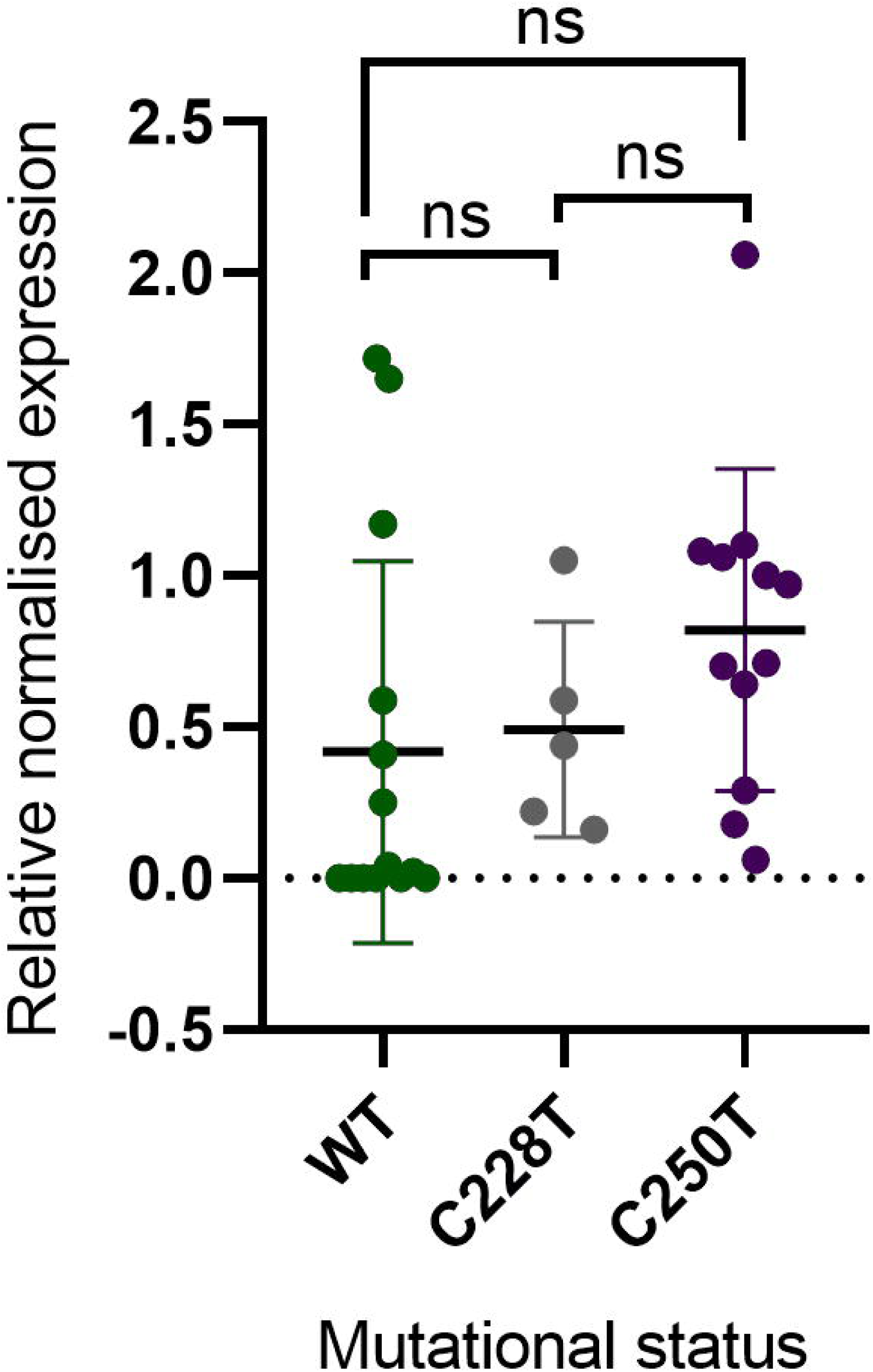
Correlation between *TERT*p mutational status and *TERT* mRNA expression in total of 31 samples.

**Fig 6.**
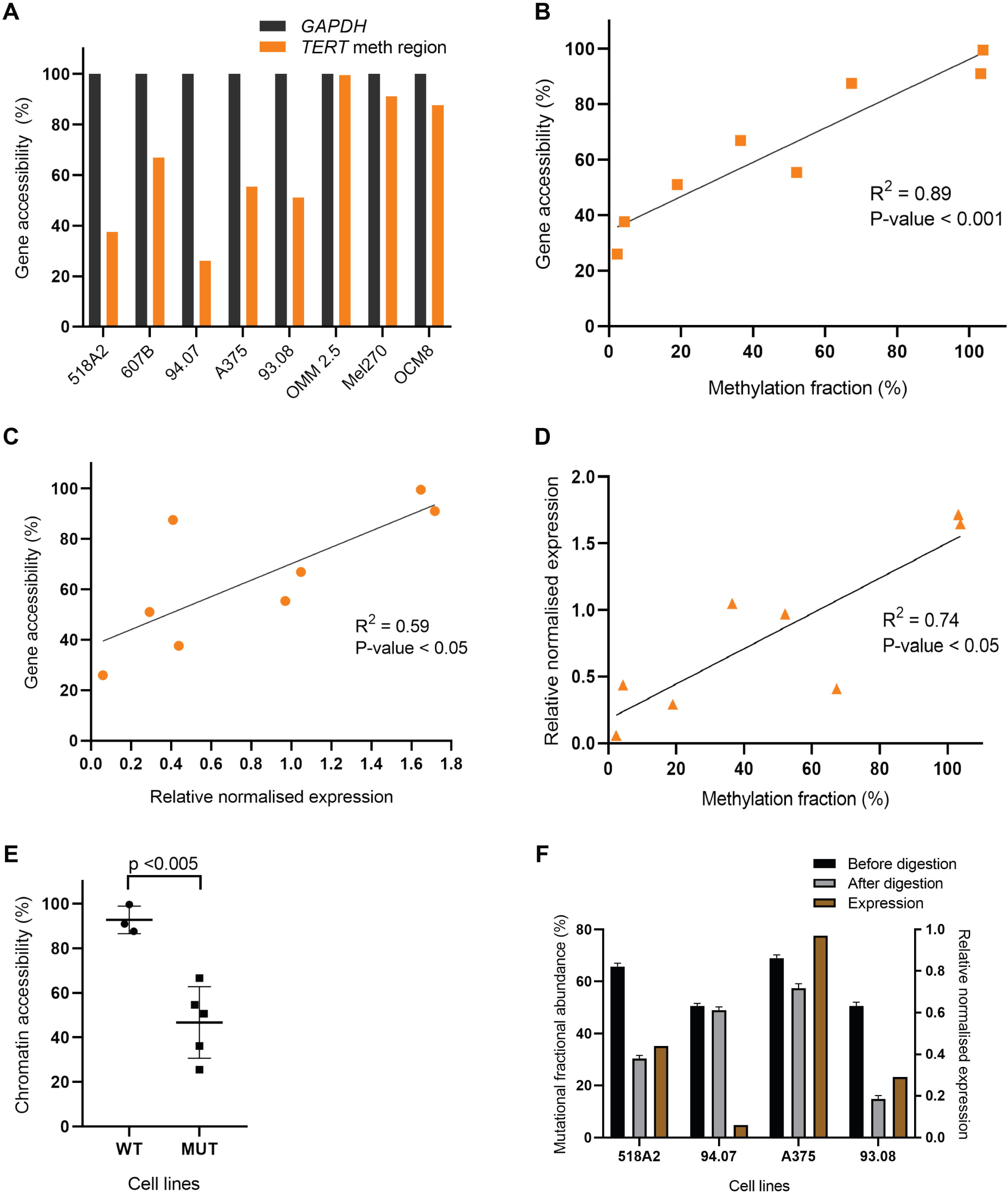
Accessibility of *TERT*p around cg11625005 in 8 melanoma cell lines. Cell lines were analysed with the EpiQ chromatin kit, and ddPCR was performed using primers and probes for positive control gene *GAPDH* and for the *TERT* methylation region, a 231-bp amplicon around cg11625005. Accessibility (%) was calculated by the ratio of the digested sample to its matched undigested sample, subtracted from 1, and subsequently normalised against the positive control *GAPDH*. **A**. Accessibility of *GAPDH* and the *TERT* methylation region, normalised against *GAPDH*. **B & C.** Correlation plots of gene accessibility around cg11625005 with the MF (%) of cg11625005 obtained through ddPCR (**B**), or with normalised expression levels via qPCR (**C**). **D**. Correlation plot between MF (%) of cg11625005 obtained through ddPCR and normalised expression levels via qPCR **E.** Comparison of WT (OMM2.5, Mel270 and OCM8) and mutated (518A2, 607B, 94.07, A375, 93.08) *TERT*-expressing cell lines subsets regarding chromatin accessibility. **F.** Mutational fractional abundance (%) in a subset of 4 *TERT*p-mutated cutaneous cell lines before and after digestion by nuclease compared to the expression.

**Fig 7.**
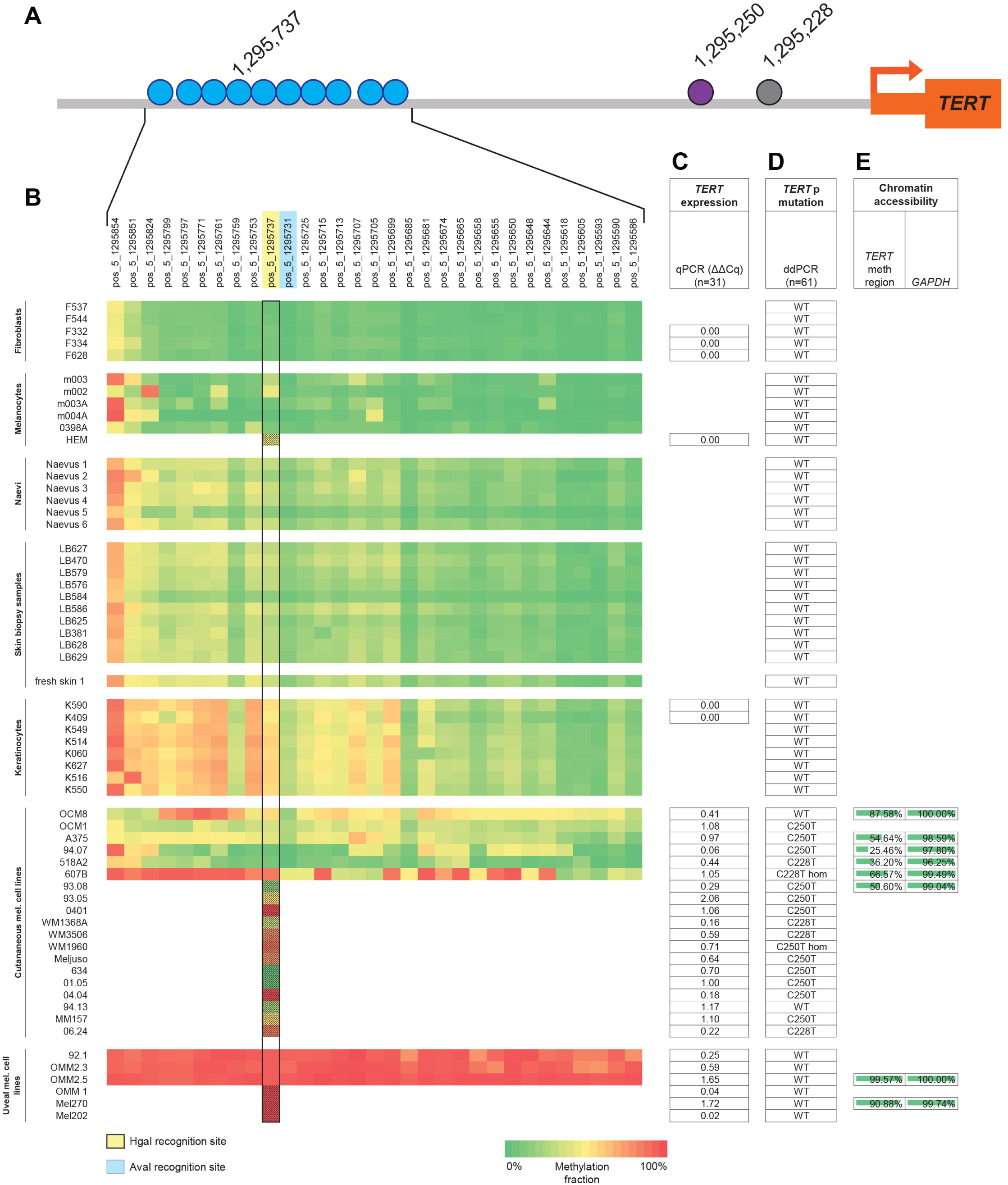
Results overview. **A**. Schematic representation of *TERT*p with the relative positions of cg11625005 (position 1,295,737 in hg19) to the *TERT*p mutations (position 1,295,228 and 1,295,250) and the transcription start site (TSS). **B**. Heat-map of methylation fraction (MF) in 31 CpG sites (top) in 44 samples (left). Yellow-marked CpG cg11625005 (position 1,295,737) is recognised by MSRE HgaI. Blue-marked CpG in 1,295,731 is recognised by MSRE AvaI. Black rectangle: MF at the cg11625005 measured either by NGS (clear squares, n=44) and by ddPCR (patterned squares, n=17; these samples were not included in the 44-sample batch subjected to NGS). **C.** *TERT* mRNA expression in 31 samples by qPCR analysed through the ΔΔCT method in Bio-Rad CFX manager software (version 3.1, Bio-Rad). **D.** *TERT*p mutations evaluated through ddPCR with commercial TERT C250T and C228T Mutation Assays in total 61 samples. **E.** Analysis of the chromatin accessibility in 8 cultured cell lines for *TERT* methylation region using *GAPDH* as a positive (constitutively expressed) control.

### Evaluation of *TERT*p mutations in a collection of skin samples and melanoma cell lines

Besides promoter methylation, somatic mutations are also known to be correlated with *TERT*p reactivation. Therefore, we characterised the *TERT*p mutational status of the sample cohort. Sanger sequencing on one naevus, fresh skin and cutaneous melanoma cell lines 518A2, 607B, A375, 94.07 and 93.08 revealed melanoma-associated *TERT* C250T and C228T mutations (Fig 4A). Aiming to use the ddPCR method to evaluate the mutational load of the samples, the *TERT* C250T and C228T mutation assays were validated in three samples of which the mutation was identified in sequencing analysis, 518A2, 607B and A375 (Fig 4B). Following the test runs, the C228T and C250T assays were used on the extended sample cohort (n=61) (S5 Table and Fig 7D). All *TERT*p-mutated samples were cutaneous melanoma cell lines, however OCM8 and 94.13 cutaneous cell lines tested wild-type. The C250T mutation was not present in combination with the C228T mutation in any sample, confirming that the mutations are mutually exclusive.

### Absence of correlation between mutational status and *TERT* expression

As the presence of mutations in the gene promoter induces *TERT* reactivation, we assessed the correlation between mutational status with *TERT* mRNA expression (n=34). When WT and mutated samples (either C228T or C250T) were compared, regardless of origin of the tissue, no significant differences for *TERT* mRNA expression were found (Fig 5). Moreover, *TERT* expression was exclusive to the melanoma cell lines, either with or without *TERT*p mutations (Fig 7C).

### *TERT* expression is correlated to chromatin accessibility

In contrast to most genes, methylation of the *TERT*p positively correlates with its mRNA expression [3, 16, 17, 19]. Although we were not able to confirm this finding, we investigated whether besides promoter methylation, other mechanisms could contribute to chromatin accessibility to transcription factors affecting *TERT*p regulation. Therefore, we analysed chromatin state in a subset of melanoma cell lines (cutaneous, 518A2, 607B, 94.07, A375, 93.08 and OCM8; and uveal, OMM2.5 and Mel270) by ddPCR methodology instead of qPCR for an accurate quantification. The positive control gene *GAPDH*, a housekeeping gene that is generally expressed in all conditions, and thus 100% accessible, was used. The accessibility in the region around cg11625005 shows a high variability, being over 90% in uveal cell lines while being intermediate to low in cutaneous melanoma cell lines (Fig 6A and an overview in Fig 7E and S6 Table). When comparing the accessibility around cg11625005 to the methylation fraction of this CpG, a significant positive correlation was observed (R^2^= 0.89, P<0.001) (Fig 6B). Another positive correlation (R^2^=0.59, P<0.05) was found when comparing the accessibility of the same region to the normalised *TERT* mRNA expression levels in these samples (Fig 6C). In actuality, in this subset of 8 cell lines, the *TERT*p methylation and gene expression show a statistically significant (P-value<0.05) positive correlation (Fig 6D). The 3 cell lines with higher MF are those with the highest chromatin accessibility (OMM2.5, Mel270 and OCM8). Remarkably, these are also the cell lines with WT-TERTp, in which the chromatin accessibility was significantly higher than in the mutated subgroup (Fig 6E).

In addition, we investigated whether the TERT accessibility originated from the mutant or the wildtype allele. For this purpose, we assessed the fractional abundance of mutated allele, in the subgroup of 4 *TERT*p-mutated cutaneous cell lines before and after nuclease digestion. 607B cell line was not included since it is homozygous for the mutation and not informative. Assuming that the nuclease digests DNA in open and accessible chromatin regions, the observed decrease in mutation fractional abundance after digestion (Fig 6F) in all 4 cell lines suggest that mutated alleles were preferably digested over WT alleles.

## Discussion

By using advanced quantification methods, we investigated the epigenetic and genetic regulation of *TERT*p in benign and malignant skin cells. Innovative ddPCR-based assays were developed and validated to assess *TERT* promoter methylation and chromatin accessibility. These methods overcome fallible bisulfite-conversion and avoid semi-quantitative qPCR and provide absolute quantification even in samples that are challenged by DNA concentration and integrity.

The methylation fraction assessed by both NGS and ddPCR was high in a variety of normal samples, of which mainly keratinocytes exceeded levels of cutaneous melanoma cell lines. This is in contrast with previous investigations on brain tumours and skin melanoma that observed a general absence of cg11625005 methylation in normal cells [3, 20]. In our study, methylation of cg11625005 at position 1,295,737 did not stand out across the CpGs in *TERT*p but seemed to be affected along with other CpG’s in the surrounding region in all samples (Fig 7B). This result suggests that context-related methylation around cg11625005 is biologically relevant in opposition to methylation of one specific CpG. Consistent with previous findings, the methylation of most samples gradually increased in the 5’ direction and decreased near the transcription start site (TSS) of the *TERT* gene (Fig 7B) [19, 25]. Regardless of the methylation status, human-derived benign cells did not express *TERT* indicating that other epigenetic mechanisms are involved (Fig 8). In contrast, analysis of tumour cell lines revealed a wide variety of promoter methylation levels (5%-100% MF). *TERT* expression was found in all tumour cell lines with or without *TERT*p mutation.

**Fig 8.**
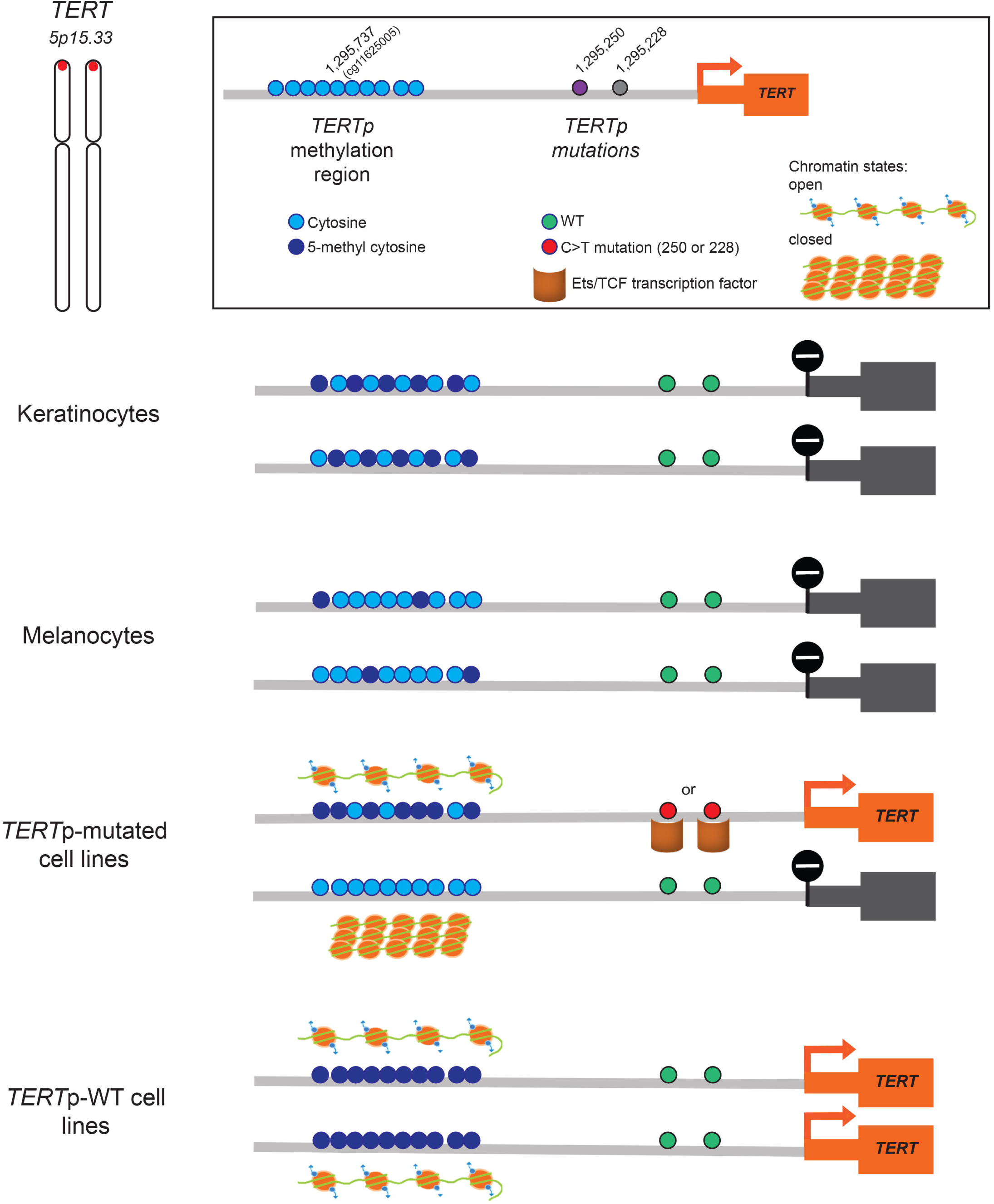
Proposed model of *TERT* transcriptional regulation. Regardless of MF at the *TERT*p methylation region, both keratinocytes and melanocytes do not show TERT expression. In *TERT*p-mutated cell lines, an intermediate MF positively correlated with chromatin accessibility, in combination with C228T/C250T *TERT* mutations allows monoallelic *TERT* expression. In *TERT*p-WT cell lines, the MF is close to 100% with a significantly higher chromatin accessibility leading to the highest expression levels.

A plethora of histone modifications result in chromatin remodelling that may change accessibility of the *TERT*p to transcription factors, such as ETS/TCF [7]. Therefore, we explored the higher-order chromatin state and its interaction with methylation levels and mRNA expression in 6 cutaneous and 2 uveal melanoma cell lines. We found that the gene accessibility around cg11625005 showed a positive correlation with the methylation and *TERT* mRNA expression in these samples.

We next investigated whether both wildtype and mutant *TERT* alleles were equally affected by higher order chromatin organization and assessed the mutational fraction upon digestion with nuclease, assuming that the nuclease only digests DNA in open and accessible chromatin regions. We could infer that, mutated alleles are more accessible, possibly favouring the binding of transcription factors, such as ETS/TCF, and consequently *TERT* expression of the mutant allele (Fig 8). The 94.07 cell line is an exception to the rule that still supports the dominant role of higher order chromatin organization since both alleles were equally resistant to nuclease digestion and presented with very low methylation fraction, explaining the lowest *TERT* expression levels among all cell lines. Our results are in line with the study from Stern *et al.* and Huang *et al.*, where the authors found that active mutant allele allows monoallelic *TERT* expression [25, 26].

Another remarkable observation in our study is that in WT *TERT*-expressing uveal melanoma cell lines, the methylation of the whole region surrounding cg11625005 is close to 100% with a significantly higher chromatin accessibility compared to *TERT*p-mutated cell lines with moderate methylation. In these cases, of *TERT*p-WT samples that show gene expression, we were not only able to confirm but also expand previous results, in which *TERT*p methylation carries out a non-canonical role, leading to transcriptional activation (Fig 8).

We conclude that ddPCR is a highly sensitive and quantifiable technique that can reliably assess methylation fractions and mutational status even in CG-rich sequences such as *TERT* gene. Further investigation in primary melanoma is needed to assess whether *TERT* methylation is predictive of worse prognosis and at which methylation fraction this phenomenon occurs [25]. Thereafter, quantification of *TERT* methylation might be used for the assessment of patient prognosis, as it is readily applicable in the clinic. Although *TERT* is one of the most affected genes in cancer, with its noncoding mutations cooperating with promoter methylation, further investigation must be conducted to fully understand all epigenetic mechanisms that collectively reactivate *TERT*.

## Material and Methods

### Samples, DNA extraction and PCR

Tissue samples were derived from anonymous patients and consisted of 11 normal skin samples, 6 frozen naevi, and low-passage cultured samples: 5 fibroblasts, 6 melanocytes and 8 keratinocytes. Primary human fibroblasts and keratinocytes were isolated from surplus human breast skin as described before [27]. Keratinocytes were used at passage 2, while fibroblasts were used at passage 3-5. The low-passage cultured fibroblasts, keratinocytes and melanocytes were a kind gift from A. El Ghalbzouri and JJ Out-Luiting [27].

We also included 19 cutaneous and 6 uveal melanoma cell lines [28]. The batch thus consisted of 39 primary skin type samples and 25 melanoma cell lines, totalling 61 samples (Table 1). The study was approved by the Leiden University Medical Center institutional ethical committee (05-036) and was conducted according to the Declaration of Helsinki Principles.

**Table 1:** Samples overview.

| Control samples |  |  |  |  | Melanoma cell lines |  |
| --- | --- | --- | --- | --- | --- | --- |
| Skin biopsy samples | Fibroblasts | Melanocytes | Keratinocytes | Naevi | Cutaneous | Uveal |
| LB627 | F537 | m003 | K590 | Naevus 1 | 0401 | OMM 2.3 |
| LB470 | F544 | m002 | K409 | Naevus 2 | WM1368A | OMM 1 |
| LB579 | F332 | m003A | K549 | Naevus 3 | 93.05 | OMM 2.5 |
| LB576 | F334 | m004A | K514 | Naevus 4 | WM3506 | Mel270 |
| LB584 | F628 | 0398A | K060 | Naevus 5 | WM1960 | Mel202 |
| LB586 |  | HEM | K627 | Naevus 6 | Meljuso | 92.1 |
| LB625 |  |  | K516 |  | 634 |  |
| LB381 |  |  | K550 |  | OCM8 |  |
| LB628 |  |  |  |  | OCM1 |  |
| LB629 |  |  |  |  | 518A2 |  |
| Fresh skin 1 |  |  |  |  | 607B |  |
|  |  |  |  |  | 94.07 |  |
|  |  |  |  |  | A375 |  |
|  |  |  |  |  | 93.08 |  |
|  |  |  |  |  | 94.13 |  |
|  |  |  |  |  | 01.05 |  |
|  |  |  |  |  | 04.04 |  |
|  |  |  |  |  | MM157 | 228 |
|  |  |  |  |  | 06.24 | 229 |

DNA was isolated using the QIAamp DNA Blood Mini Kit and the DNeasy Blood & Tissue Kit (both from Qiagen, Hilden, Germany).

Conventional PCR was performed using the PCR-sequencing kit (Thermo Fisher Scientific, Waltham, MA, USA), containing 10X reaction buffer, MgCl_2_ (50mM), dNTP mix (10nM, Fermentas/Thermo Fisher Scientific), primer mix (900nM each), PlatinumX Taq enzyme (2.5U), 50ng DNA and Aqua B. Braun RNase-free water. A PCR for CG-rich sequences was performed on 50ng DNA using the PCRX Enhancer System (Thermo Fisher Scientific), containing 10X PCRX amplification buffer, MgSO_4_ (50mM), dNTP mix (10nM), primer mix (900nM each), PlatinumX Taq enzyme (2.5U) and Aqua B. Braun RNase-free water. The samples were amplified in C1000 Touch Thermal Cycler (Bio-Rad Laboratories, Inc., Hercules, CA, USA).

### Promoter methylation determination

#### Bisulfite conversion and next-generation sequencing (NGS)-based deep bisulfite sequencing

DNA was bisulfite-converted (BC) using the EZ DNA Methylation™ Kit (Zymo Research, Irvine, CA, USA) according to the manufacturer protocol (version 1.2.2). BC samples were amplified using the PCRX Enhancer System in the program: 1 cycle of 95°C for 3 minutes, 8 cycles of 95°C for 30 seconds, 58°C for 30 seconds, reducing 1°C/cycle, and 68°C for 1 minute, then 36 cycles of 95°C and 53°C for 30 seconds each, and 68°C for 1 minute, followed by 1 cycle of 68°C for 3 minutes. Tailed primers were used for amplification (900nM each; S1 Table). Samples were sequenced through next-generation sequencing (NGS), MiSeq, 2×300bp paired-end, at Leiden Genome Technology Centre (LGTC).

#### Novel design of a ddPCR assay using methylation-sensitive restriction enzymes (MSREs) to determine *TERT*p methylation fraction

The methylation fraction (MF) of the CpG (cg11625005) in position 1,295,737 was determined by an in-house designed ddPCR assay in combination with HgaI methylation-sensitive restriction enzyme (MSRE) that cleaves this CpG when unmethylated, as described by Nell *et al.* [24]. 100ng DNA sample was incubated with HgaI (2U/μl) and appurtenant 10X NEBuffer 1.1 (both from New England Biolabs, Bioké, Leiden, The Netherlands) for 60 minutes at 37°C and 65°C for 20 minutes. To assess the MF of a CpG adjacent to cg11625005, located in 1,295,731, the MSRE AvaI (10U/μl; New England Biolabs) was employed, which recognises this CpG and cleaves it when unmethylated. Incubation of the DNA samples with AvaI was performed with 10X CutSmart buffer for 15 minutes at 37°C and subsequently 65°C for 20 minutes. For ddPCR reaction, 60ng DNA digested or undigested by HgaI, 2x ddPCR SuperMix for Probes (no dUTP), primers (900nM each), a FAM-labelled in-house-designed probe for the CpG site of interest (250nM, Sigma, St. Louis, MO, USA), and 20X HEX-labelled CNV *TERT* reference primer/probe (Bio-Rad) for total *TERT* amplicon count. The primer and probe sequences are presented in S2 Table. The amplification protocol used: 1 cycle of 95°C for 10 minutes, 40 cycles of 94°C for 30 seconds and 60°C for 1 minutes, and 1 cycle of 98°C for 10 minutes, all at ramp rate 2°C/s. Droplets were analysed through a QX200 droplet reader (Bio-Rad) using QuantaSoft software version 1.7.4 (Bio-Rad). Raw data was uploaded in online digital PCR management and analysis application Roodcom WebAnalysis (version 1.4.2, https://www.roodcom.nl/webanalysis/) [24], in which the MF was calculated by dividing the CNV of the digested sample with that of the paired undigested sample.

### Assessment of mutational status

#### Sanger sequencing

The presence of the C228T and C250T *TERT*p mutations in some samples was evaluated by conventional Sanger sequencing. DNA samples were amplified through the PCRX Enhancer System (Thermo Fisher Scientific) using primers (Sigma-Aldrich) and amplification program described by McEvoy *et al.* [29].

#### Mutation analysis using commercial TERT C250T and C228T mutation assays

For most of the samples, the *TERT*p mutations were detected by the ddPCR technique according to protocol described by Corless *et al.* [30], using the *TERT* C250T_113 Assay and C228T_113 Assay (unique assay ID dHsaEXD46675715 and dHsaEXD72405942, respectively; Bio-Rad). Both assays include FAM-labelled probes for the C250T and C228T mutations respectively, HEX-labelled wild-type (WT) probes, and primers for a 113-bp amplicon that encompasses the mutational sites. The ddPCR reaction mix comprised 1X ddPCR Supermix for Probes (No dUTP), Betaine (0.5M; 5M stock), EDTA (80mM; 0.5M stock, pH 8.0, Thermo Fisher Scientific), CviQI restriction enzyme (RE; 2.5U; 10U/μl stock, New England BioLabs), the *TERT* assay, and 50ng DNA. Droplets were generated in QX200 AutoDG system (Bio-Rad) and amplified in T100 Thermal Cycler (Bio-Rad) according to the recommended cycling conditions and analysed through a QX200 droplet reader (Bio-Rad) using QuantaSoft software version 1.7.4.0917 (Bio-Rad).

### Chromatin accessibility

#### Cell culture and treatment to assess chromatin states

Cutaneous melanoma cell lines A375, 518A2, 607B, 94.07, 93.08, OMM2.5, Mel270 and OCM8 were cultured for 22 days in 9-cm Cellstar® cell culture dishes (Greiner Bio-One GmbH, Frickenhausen, Germany) with Dulbecco’s modified eagle medium (DMEM; Sigma-Aldrich) supplemented with 10% FCS, Penicillin (100U/ml), and Streptomycin (100μg/ml; both from Lonza, Verviers, Belgium) until roughly 95% confluent. Then, different densities (10,000, 20,000, 40,000 and 80,000 cells) of the above-mentioned cell lines were seeded in duplicate into a 48-well plate (Corning Costar, Sigma-Aldrich) required for the EpiQ chromatin assay. The EpiQ™ Chromatin Analysis Kit (Bio-Rad) was performed according to manufacturer’s instructions. Briefly, after 2 days each cell line was 85%-95% confluent. The cells were permeabilised and treated with EpiQ chromatin digestion buffer with or without nuclease for 1 hour at 37°C. Following incubation with EpiQ stop buffer for 10 minutes at 37°C, the DNA samples were purified using alcohol and DNA low- and high-stringency wash solutions. The genomic DNA was eluted in DNA elution solution.

#### Novel design of a ddPCR assay to assess chromatin opening state

The analysis was performed using ddPCR rather than qPCR, to achieve quantifiable results using *GAPDH* expression as positive control. The reaction mix consisted of 2x ddPCR Supermix for Probes (No dUTP, Bio-Rad), 20x HEX-labelled CNV *TERT* reference primer/probe (Bio-Rad), 50ng DNA, and primers (900nM each) and FAM-labelled probes (250nM) for *GAPDH*, or the methylation region around cg11625005 (S3 Table). Samples were amplified according to the program of the CNV *TERT* reference primer/probe as described. Gene accessibility was quantified by the digestion fraction between the digested and undigested samples, subtracted from 1.

#### RNA isolation, cDNA synthesis and quantitative real-time PCR

RNA was obtained using the FavorPrep Tissue Total RNA Extraction Mini Kit (Favorgen Biotech, Vienna, Austria) according to manufacturer’s instructions for animal cells. cDNA was synthesised through the iScript™ cDNA Synthesis Kit (Bio-Rad) according to recommended protocol. *TERT* mRNA expression was assessed by qPCR performed with 3.5ng DNA, IQ SYBR Green Supermix (2x; Bio-Rad), and 0.5μM PCR primers (Sigma-Aldrich; S4 Table) in a Real-Time PCR Detection System CFX96 (Bio-Rad) and normalised to reference gene expression (*RPS11, TBP* and *CPSF6*, S4 Table). Data was analysed through the ΔΔCT method in Bio-Rad CFX manager software (version 3.1, Bio-Rad).

### Statistical Analysis

MF obtained using ddPCR was calculated with 95% confidence interval through RoodCom WebAnalysis (version 1.4.2). Significant testing of linear regression and multiple comparisons in correlation plots was performed through GraphPad Prism (version 8 for Windows, GraphPad Software, CA, USA).

## Acknowledgements

We thank Mieke Versluis, Wim Zoutman, AG Jochemsen and Mijke Visser for useful discussions. We would like to thank Coby Out and Tim van Groningen for the assistance with cell culturing.

## Supporting information

**S1 Table.** Tailed primers used for amplification of 325-bp region in bisulfite-converted samples.

**S2 Table.** Primers and probe sequences to amplify the 106-bp amplicon in a novel design of a ddPCR assay to determine the methylation fraction.

**S3 Table.** Primers and probe sequences to amplify the 231-bp region encompassing 31 CpG sites around the cg11625005 in a novel ddPCR assay to assess the chromatin state.

**S4 Table.** Primer and probe sequences for *TERT* expression in qPCR.

**S5 Table.** Overview of the methylation fraction (measured by ddPCR and NGS), mutational status and *TERT* mRNA expression of our sample cohort (n=61).

**S6 Table.** Overview of the methylation fraction (measured by ddPCR and NGS), mutational status and *TERT* mRNA expression and chromatin accessibility in the subset of melanoma cell lines present of our cohort (n=25).

