## Supporting information for "Interplay between *TERT* promoter mutations and methylation culminates in chromatin accessibility and *TERT* expression"

**S1 Table.** Tailed primers used for amplification of 325-bp region in bisulfite-converted samples.

| | |
|---|---|
| Forward primer (5'-3') | [GATGTGTATAAGAGACAG]AGGGGTTATGATGTGGAGGT |
| Reverse primer (5'-3') | [CGTGTGCTCTTCCGATCT]TTACTCATAATAAAAACCCCTC |

Note: Primer tail between square brackets

**S2 Table.** Primers and probe sequences to amplify the 106-bp amplicon in a novel design of a ddPCR assay to determine the methylation fraction.

| | |
|---|---|
| Forward primer (5'-3') | GTGAAGGGGAGGACGGAGG |
| Reverse primer (5'-3') | GTGTTGCAGGGAGGCACT |
| Probe (5'-3') | TAGACGCGGCTGGGGACGAA |

**S3 Table.** Primers and probe sequences to amplify the 231-bp region encompassing 31 CpG sites around the cg11625005 in a novel ddPCR assay to assess the chromatin state.

| | Forward primer (5'-3') | Reverse primer (5'-3') | Probe (5'-3') |
|---|---|---|---|
| *GAPDH* | CCTTGACTCCCTAGTGTCCT | ATTTATAGAAACCGGGGGCG | CGGGGCCCACACGCTCGGT |
| *TERT* methylation region | GCCTAGGCTGTGGGGTAAC | CCCGTCCAGGGAGCAA | GCGGCGACCCTTTGGCCGC |

**S4 Table.** Primer and probe sequences for *TERT* expression in qPCR.

| | | Forward primer (5'->3') | Reverse primer (5'->3') |
|---|---|---|---|
| *TERT* | ex9-10 | ATCCTCTCCACGCTGCTCT | CCAACAAGAAATCATCCACCA |
| Ref. | *RPS11* | AACATCGGTCTGGGCTTC | AGTGAAGGGGCATTTCTTGT |
| | *TBP* | CACGAACCACGGCACTGATT | TTTTCTTGCTGCCAGTCTGGAC |
| | *CPSF6* | AAGATTGCCTTCATGGAATTGAG | TCGTGATCTACTATGGTCCCTCTCT |

Ref.: reference (household)

**S5 Table.** Overview of the methylation fraction (measured by ddPCR and NGS), mutational status and *TERT* mRNA expression of our sample cohort (n=61).

| | Type of samples | Sample ID | Methylation fraction (cg11625005) | | *TERT* p mutation | *TERT* expression |
|---|---|---|---|---|---|---|
| | | | ddPCR (n=59) | NGS (n=44) | ddPCR (n=61) | qPCR (ΔΔCq) (n=31) |
| **1** | **Uveal cell lines** | OMM 2.3 | 103.8% | 96% | WT | 0.59 |
| **2** | | OMM 1 | 103.4% | | WT | 0.04 |
| **3** | | OMM 2.5 | 103.8% | 97% | WT | 1.65 |
| **4** | | Mel270 | 103.2% | | WT | 1.72 |
| **5** | | Mel202 | 107.8% | | WT | 0.02 |
| **6** | | 92.1 | 100.4% | 96% | WT | 0.25 |
| **7** | **Cutaneous cell lines** | 0401 | 100.0% | | C250T | 1.06 |
| **8** | | WM1368A | 35.9% | | C228T | 0.16 |
| **9** | | 93.05 | 53.4% | | C250T | 2.06 |
| **10** | | WM3506 | 78.8% | | C228T | 0.59 |
| **11** | | WM1960 | 87.3% | | C250T hom | 0.71 |
| **12** | | Meljuso | 74.4% | | C250T | 0.64 |
| **13** | | 634 | 4.4% | | C250T | 0.70 |
| **14** | | OCM8 | 67.3% | 56% | WT | 0.41 |
| **15** | | OCM1 | 36.2% | 37% | C250T | 1.08 |
| **16** | | 518A2 | 4.3% | 8% | C228T | 0.44 |
| **17** | | 607B | 36.5% | 86% | C228T hom | 1.05 |
| **18** | | 94.07 | 2.3% | 5% | C250T | 0.06 |
| **19** | | A375 | 52.4% | 41% | C250T | 0.97 |
| **20** | | 93.08 | 19.0% | | C250T | 0.29 |
| **21** | | 01.05 | 7.5% | | C250T | 1.00 |
| **22** | | 04.04 | 102.6% | | C250T | 0.18 |
| **23** | | 94.13 | 23.7% | | WT | 1.17 |
| **24** | | MM157 | 55.1% | | C250T | 1.10 |
| **25** | | 06.24 | 83.0% | | C228T | 0.22 |
| **26** | **Skin biopsy samples** | LB627 | 21.9% | 29% | WT | |
| **27** | | LB470 | 26.0% | 36% | WT | |
| **28** | | LB579 | 23.4% | 30% | WT | |
| **29** | | LB576 | 18.3% | 30% | WT | |
| **30** | | LB584 | 10.1% | 15% | WT | |
| **31** | | LB586 | 31.0% | 34% | WT | |
| **32** | | LB625 | | 25% | WT | |
| **33** | | LB381 | 26.3% | 29% | WT | |
| **34** | | LB628 | 22.7% | 28% | WT | |
| **35** | | LB629 | 24.3% | 28% | WT | |
| **36** | | fresh skin 1 | 25.8% | 33% | WT | |
| **37** | **Keratinocytes** | K590 | 41.5% | 54% | WT | 0.00 |
| **38** | | K409 | 49.4% | 49% | WT | 0.00 |
| **39** | | K549 | 34.0% | 56% | WT | |
| **40** | | K514 | 43.5% | 59% | WT | |
| **41** | | K060 | 41.4% | 56% | WT | |
| **42** | | K627 | 42.0% | 59% | WT | |
| **43** | | K516 | 41.1% | 59% | WT | |
| **44** | | K550 | 34.1% | 56% | WT | |
| **45** | **Melanocytes** | m003 | 9.5% | 29% | WT | |
| **46** | | m002 | 7.1% | 44% | WT | |
| **47** | | m003A | | 7% | WT | |
| **48** | | m004A | 3.6% | 2% | WT | |
| **49** | | 0398A | 4.3% | 9% | WT | |
| **50** | | HEM | 43.9% | | WT | 0.00 |
| **51** | **Fibroblasts** | F537 | 3.7% | 5% | WT | |
| **52** | | F544 | 4.0% | 7% | WT | |
| **53** | | F332 | 3.5% | 9% | WT | 0.00 |
| **54** | | F334 | 4.4% | 8% | WT | 0.00 |
| **55** | | F628 | 3.3% | 8% | WT | 0.00 |
| **56** | **Naevi** | Naevus 1 | 26.4% | 31% | WT | |
| **57** | | Naevus 2 | 13.8% | 20% | WT | |
| **58** | | Naevus 3 | 26.9% | 33% | WT | |
| **59** | | Naevus 4 | 25.8% | 31% | WT | |
| **60** | | Naevus 5 | 10.2% | 15% | WT | |
| **61** | | Naevus 6 | 23.1% | 33% | WT | |

**S6 Table.** Overview of the methylation fraction (measured by ddPCR and NGS), mutational status and *TERT* mRNA expression and chromatin accessibility in the subset of melanoma cell lines present of our cohort (n=25).

| | **Type of samples** | **Sample ID** | **Methylation fraction (cg11625005)** | | ***TERT* p mutation** | ***TERT* expression** | **Gene accessibility** | |
|---|---|---|---|---|---|---|---|---|
| | | | ddPCR | NGS | ddPCR | qPCR (ΔΔCq) | *TERT* meth site | *GAPDH* |
| **1** | **Uveal cell lines** | OMM 2.3 | 103.8% | 96% | WT | 0.59 | | |
| **2** | | OMM 1 | 103.4% | | WT | 0.04 | | |
| **3** | | OMM 2.5 | 103.8% | 97% | WT | 1.65 | 99.6% | 100.0% |
| **4** | | Mel270 | 103.2% | | WT | 1.72 | 90.9% | 99.7% |
| **5** | | Mel202 | 107.8% | | WT | 0.02 | | |
| **6** | | 92.1 | 100.4% | 96% | WT | 0.25 | | |
| **7** | **Cutaneous cell lines** | 0401 | 100.0% | | C250T | 1.06 | | |
| **8** | | WM1368A | 35.9% | | C228T | 0.16 | | |
| **9** | | 93.05 | 53.4% | | C250T | 2.06 | | |
| **10** | | WM3506 | 78.8% | | C228T | 0.59 | | |
| **11** | | WM1960 | 87.3% | | C250T hom | 0.71 | | |
| **12** | | Meljuso | 74.4% | | C250T | 0.64 | | |
| **13** | | 634 | 4.4% | | C250T | 0.70 | | |
| **14** | | OCM8 | 67.3% | 56% | WT | 0.41 | 87.6% | 100.0% |
| **15** | | OCM1 | 36.2% | 37% | C250T | 1.08 | | |
| **16** | | 518A2 | 4.3% | 8% | C228T | 0.44 | 36.2% | 96.3% |
| **17** | | 607B | 36.5% | 86% | C228T hom | 1.05 | 66.6% | 99.5% |
| **18** | | 94.07 | 2.3% | 5% | C250T | 0.06 | 25.5% | 97.8% |
| **19** | | A375 | 52.4% | 41% | C250T | 0.97 | 54.6% | 98.6% |
| **20** | | 93.08 | 19.0% | | C250T | 0.29 | 50.6% | 99.0% |
| **21** | | 01.05 | 7.5% | | C250T | 1.00 | | |
| **22** | | 04.04 | 102.6% | | C250T | 0.18 | | |
| **23** | | 94.13 | 23.7% | | WT | 1.17 | | |
| **24** | | MM157 | 55.1% | | C250T | 1.10 | | |
| **25** | | 06.24 | 83.0% | | C228T | 0.22 | | |
